# Clomipramine inhibits dynamin GTPase activity by L-α-phosphatidyl-L-serine stimulation

**DOI:** 10.1101/2022.09.09.507370

**Authors:** Hiroshi Miyoshi, Masahiro Otomo, Kiyofumi Takahashi

**Author notes:** Corresponding author: Department of Microbiology, St. Marianna University School of Medicine, 2-16-1 Sugao, Miyamae, Kawasaki 216-8511, Japan. Shinkohkai izukan-nami hospital, 1694 Hirai, Kan-nami, Tagata, Shizuoka 419-0107 Japan (Masahiro Otomo). Research Organization for Nano and Life Innovation, Waseda University, Tokyo, Japan (Kiyofumi Takahashi). All authors contributed equally.

## Abstract

Three dynamin isoforms play critical roles in clathrin-dependent endocytosis. Severe acute respiratory syndrome coronavirus 2 (SARS-CoV-2) enters host cells *via* clathrin-dependent endocytosis. We previously reported that 3-(3-chloro-10,11-dihydro-5*H*-dibenzo[*b*,*f*]azepin-5-yl)-*N*,*N*-dimethylpropan-1-amine (clomipramine) inhibits the GTPase activity of dynamin 1, which is in mainly neuron. Therefore, we investigated whether clomipramine inhibits the activity of other dynamin isoforms in this study. We found that, similar to its inhibitory effect on dynamin 1, clomipramine inhibited the L-α-phosphatidyl-L-serine-stimulated GTPase activity of dynamin 2, which is expressed ubiquitously, and dynamin 3, which is expressed in the lung. Inhibition of GTPase activity raises the possibility that clomipramine can suppress SARS-CoV-2 entry into host cells.

**Graphical Abstract:** 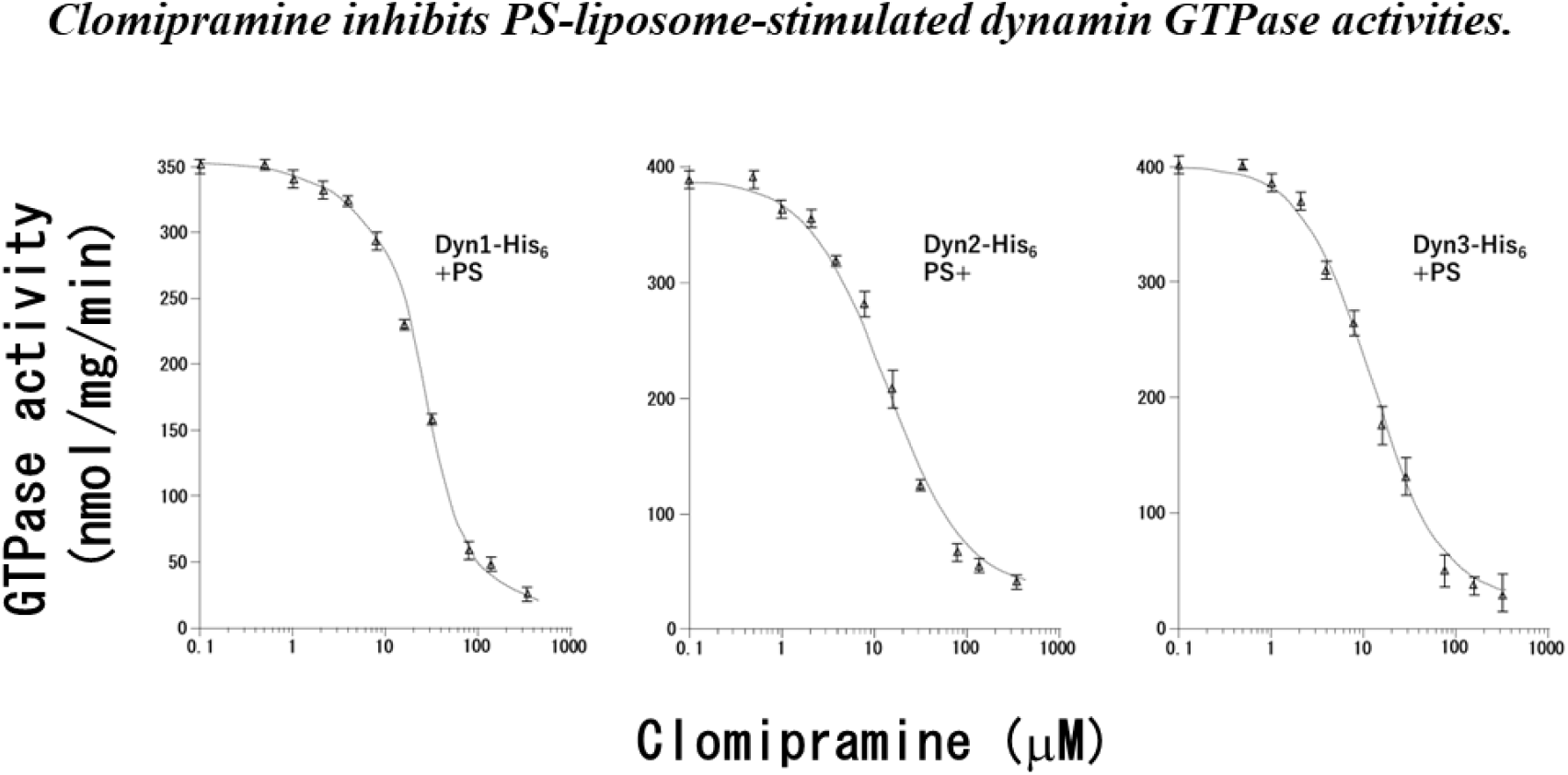

## Introduction

Dynamin (Dyn) GTPase plays a critical role in clathrin- and caveolae-dependent endocytosis (*1–3*). Mammals express three Dyn isoforms. Although Dyn 2 is expressed ubiquitously, Dyn 1 is assumed to be neuron-specific. Dyn 3 is expressed mainly in the testis, brain, and lungs (*4*).

Dyn GTPase activity’s essential function is producing mechanical force to drive membrane fission during vesicle budding during endocytosis (*1, 5*). Notably, endocytosis driven by Dyn GTPase activity is involved in the intracellular transportation of transferrin (*6*).

All Dyn isoforms carry four functional domains: a GTPase domain, pleckstrin homology domain, GTPase effector domain, and proline/arginine-rich domain (*1*). These functional domains act in concert during vesicle budding (*7*), and lipid species such as L-α-phosphatidyl-L-serine (PS) potentiate dynamin activity (*8*). Dyn GTPase activities are inhibited by cationic surface-active compounds such as myristyl trimethyl ammonium bromide (MiTMAB). MiTMAB competes with PS for binding to the pleckstrin homology domain in Dyn, but it does not compete with GTP to bind Dyn (*9*).

Cells can internalize many resources from the external cell environment *via* endocytosis, and non-enveloped viruses (e.g., hepatitis C virus) generally leverage related mechanisms to enter a cell following the same endocytic pathway as resources (*10*). Many enveloped viruses, such as severe acute respiratory syndrome coronavirus 2 (SARS-CoV-2), enter cells *via* endocytosis. The SARS-CoV-2 spike glycoprotein binds to host angiotensin-converting enzyme 2 (ACE2), thereby mediating membrane fusion and enabling virus entry into cells (*11, 12*).

Recently, Kato *et al*. reported that the tricyclic antidepressant clomipramine suppressed ACE2-mediated SARS-CoV-2 entry into host cells (*13*). Basic research has used Clomipramine as a reagent to inhibit endocytosis (*14*).

We previously reported that several antipsychotic treatments inhibited the GTPase activity of Dyn 1 and Dyn 2 and that the suppression of dynamin GTPase activity by those antipsychotic drugs led to the inhibition of dynamin-driven endocytosis. Moreover, we explained that (1*S*)-*cis*-4-(3,4-dichlorophenyl)-1,2,3,4-tetrahydro-*N*-methyl-1-naphthalenamine (sertraline) is a competitive inhibitor of dynamin I GTPase concerning both GTP and PS. (*15, 16*). In a previous study, we also reported that clomipramine inhibited the GTPase activity of neuron-specific Dyn 1 (*15*). However, the inhibition of the activity of lung-specific Dyn 3 and the activity of ubiquitous Dyn 2 by clomipramine has not yet been investigated. Therefore, the inhibition of Dyn isoform GTPase activity by clomipramine was measured to understand the inhibitory mechanism.

## Materials and Methods

### Materials

The following materials were obtained from the cited sources and used in this study: Pfu Ultra II Fusion HS DNA Polymerase (Agilent Technologies); restriction enzymes and DNA ligation kit ver.2 (Takara Bio); pET21a and *E. coli* Rosetta2 (DE3) (Merck); TALON Metal Affinity Resin (Clontech); Mono Q 5/50 GL (GE Healthcare); L-α-phosphatidyl-L-serine (Sigma Aldrich); clomipramine (Sigma); and HRP–conjugated anti Penta-His antibody (Qiagen). Other chemicals used in this study were of an analytical grade or higher. Clomipramine, sertraline, and MiTMAB were prepared in stock solutions consisting of 50% (v/v) DMSO and 30 mM Tris-HCl pH 7.4 and diluted in 30 mM Tris-HCl, pH 7.4, before use in the assay.

### Plasmid construction, protein expression and purification of Dyn isoforms

All Dyn isoform genes were amplified by PCR as described previously (*15, 16*). These Dyn genes were cloned into separate pET21a plasmids to create C-terminal-His_6_-tagged Dyn isoform protein (Dyns-His_6_) vectors. We expressed Dyns-His_6_ from *E. coli* Rosetta2 (DE3) by transforming with those Dyn isoforms-coding pET21a expression plasmids. As previously reported, the expressed Dyns-His6 were purified to homogeneity on a MonoQ column after TALON Metal Affinity Resin purification (*15–17*). The purified proteins were confirmed by 15 % SDS-PAGE.

### Preparation of PS-liposomes and performance of a GTPase assay with purified Dyns-His6

Forty microlitres of PS (10 mg ml^−1^) in chloroform/methanol (95:5) was evaporated to a final 5-μl volume, and then PS was suspended in 1 ml of 30 mM Tris-HCl, pH 7.4, and sonicated for 2 min on ice to yield a working solution of 400 μg ml^−1^. GTPase assays with the purified Dyns-His_6_ were performed *via* sensitive malachite green colorimetric detection of orthophosphate (Pi), as described previously (*15–17*). Phosphate release was quantified and the levels were compared against a standard curve of sodium dihydrogen orthophosphate monohydrate after each experiment. KaleidaGraph 4.0 (Synergy Software) was used to plot and analyze curves using non-linear regression.

### Blue native (BN)-PAGE

BN-PAGE was performed as previously reported (*17*). Purified Dyn2-His_6_ (2 μM) were incubated at room temperature for 30 min with 2 mM PS-liposomes in TBS, 2 mM PS-liposomes + 1% SDS in TBS, 1 mM clomipramine + 2 mM PS-liposomes in TBS, or 1 mM clomipramine + 2 mM PS-liposomes + 1% SDS in TBS, respectively. The incubation solutions were supplemented with BN-PAGE sample buffer (5% Coomassie Brilliant Blue G-250; 500 mM 6-aminocaproic acid; 100 mM Bis–Tris-HCl, pH 7.0; and 1 mM phenylmethylsulfonyl fluoride) and incubated on ice for 5 min. All samples were resolved in 4 to 16% gels by BN-PAGE and measured by western blotting with HRP–conjugated anti-Penta-His antibody.

## Results

### Dyns-His_6_ was activated by PS-liposomes

We expressed and purified Dyns-His_6_. Those purified Dyns-His_6_ were confirmed by 15 % SDS-PAGE as shown in Fig. 1A. To confirm the GTPase activities of purified Dyns-His_6_, malachite green colorimetric detection of released orthophosphate was performed to confirm the GTPase activity levels of purified Dyns-His_6_ with/without PS-liposome treatment. Purified Dyns-His_6_ at all concentrations hydrolyzed GTP to release Pi in a time-dependent manner, and the GTPase activity was stimulated by the PS-liposomes (Fig. 1B and C). The addition of PS-liposomes increased Dyn1-His_6_ activity by 5.1-fold, Dyn2-His_6_ activity by 4.7-fold, and Dyn3-His_6_ activity by 4.0-fold (Fig. 1B *versus* C, 60 min).

**Fig. 1.**
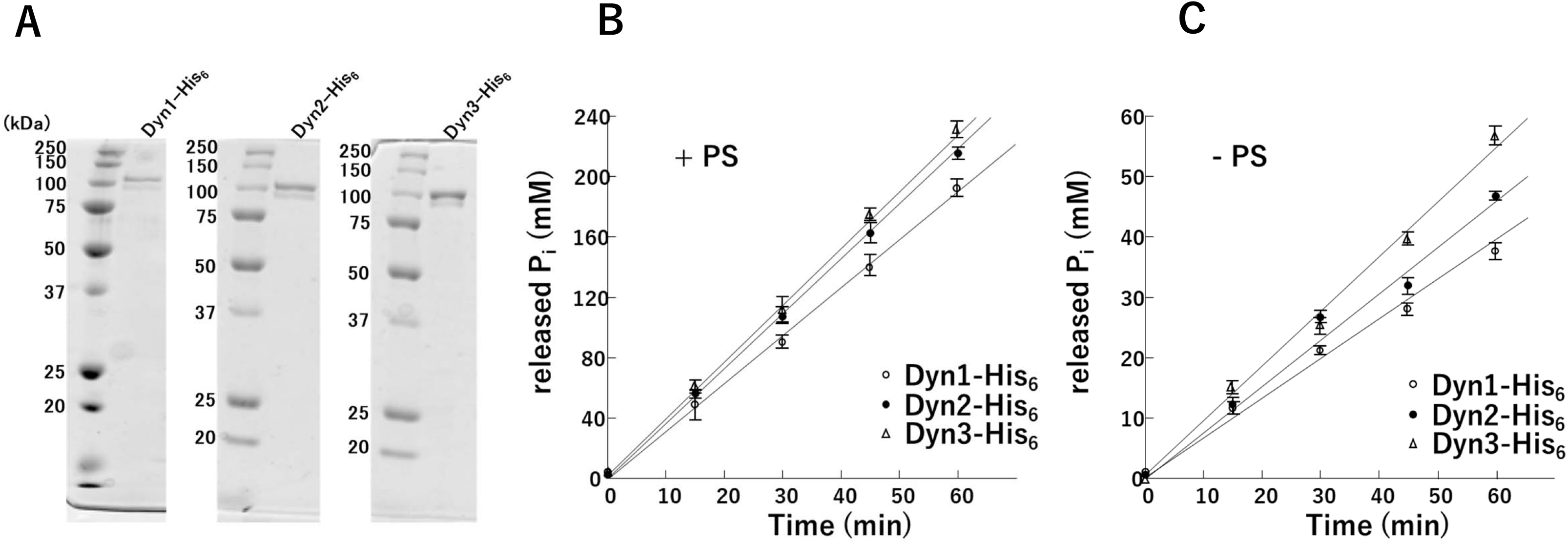
Dyns-His6 was activated by PS-liposomes. (A) Purified Dyns-His_6_ in this study. The purified proteins were resolved by 15 % SDS-PAGE and stained with Coomassie Brilliant Blue. (B) The GTPase activities of Dyn1-His_6_ (hollow circles), Dyn2-His_6_ (solid circles), and Dyn3-His_6_ (triangles) were measured as the concentration of released Pi. GTPase activities of Dyns-His_6_ isomers in the presence of 5.0 μM PS-liposomes. The GTPase activities of 20 nM Dyns-His_6_ isoforms were measured at 30 °C. The concentration of Pi released is plotted as a function of time. (C) GTPase activities of 20 nM Dyns-His_6_ isomers in the absence of PS-liposomes. All Dyn-His_6_ isoforms released orthophosphate *via* GTP hydrolysis in a time-dependent manner.

### Clomipramine inhibits the GTPase activity of all dynamin isoforms in the presence of PS-liposomes

GTPase assay was performed to evaluate the inhibition of Dyns-His_6_ GTPase activity by clomipramine, and the activity levels were compared with those of sertraline and MiTMAB (Fig. 2 and Fig. 3). In the presence of PS-liposomes, clomipramine showed significant inhibition of GTPase activity (Fig. 2). The IC_50_ values for Dyn1-His_6_ were 5.5 ± 1.2 μM (sertraline), 8.1 ± 3.7 μM (MiTMAB) and 22.0 ± 2.2 μM (clomipramine) in the presence of PS-liposomes (Fig. 2A). For Dyn2-His_6_, the IC_50_ values were 4.9 ± 0.6 μM (sertraline), 9.2 ± 2.5 μM (MiTMAB) and 12.8 ± 3.2 μM (clomipramine) in the presence of PS-liposomes (Fig. 2B). The IC_50_ values for Dyn3-His_6_ were 4.4 ± 0.5 μM (sertraline), 7.8 ± 3.0 μM (MiTMAB) and 11.3 ± 2.9 μM (clomipramine) in the presence of PS-liposomes (Fig. 2C). In contrast to MiTMAB, clomipramine showed no overt inhibition of GTPase activity, similar to sertraline, in the absence of PS-liposomes (Fig. 3).

**Fig. 2.**
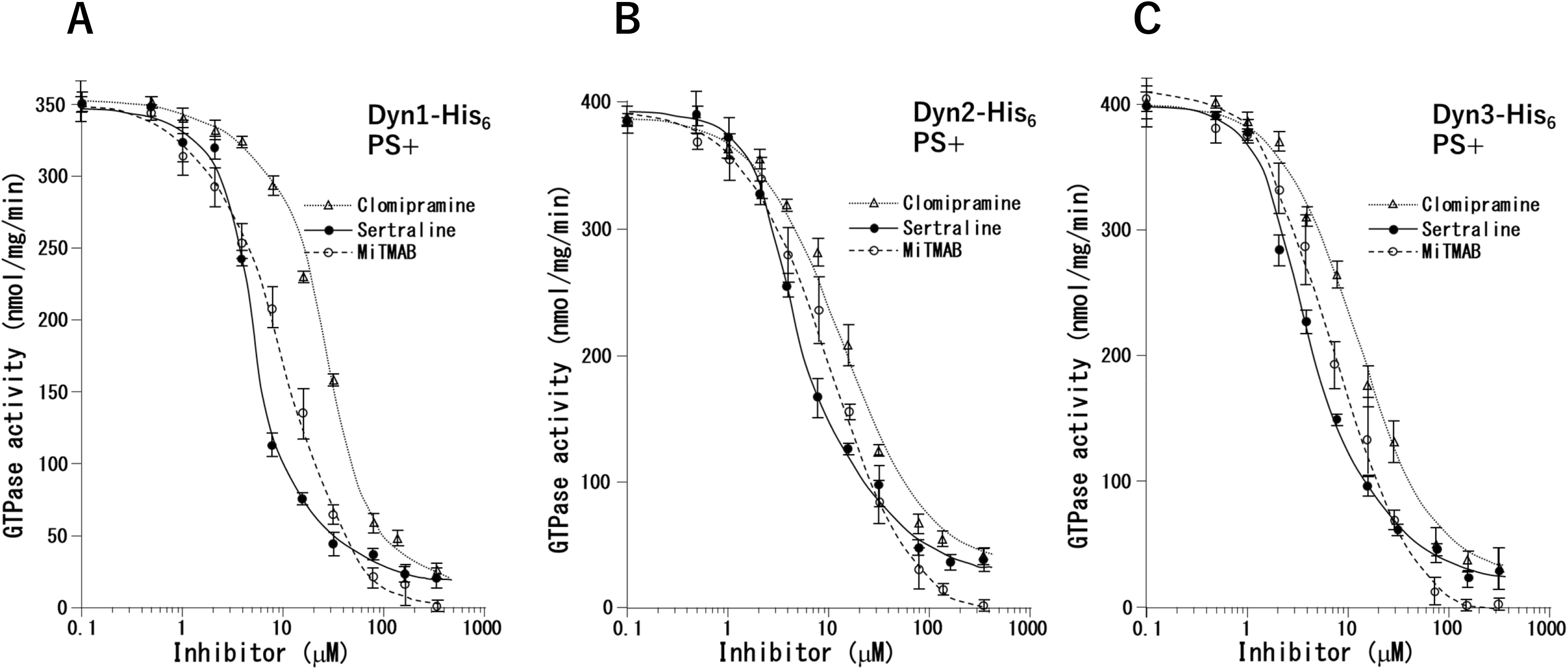
Clomipramine inhibits dynamin GTPase activity by PS-liposome-stimulation. The GTPase activity of purified Dyns-His_6_ (20 nM) was determined in the presence of various concentrations of MiTMAB (hollow circles), sertraline (solid circles), or clomipramine (triangles). All results are representative of at least 3 independent experiments. (A) GTPase activity of Dyn1-His_6_ in the presence of 5.0 μM PS-liposomes. (B) GTPase activity of Dyn2-His_6_ in the presence of 5.0 μM PS-liposomes. (C) GTPase activity of Dyn3-His_6_ in the presence of 5.0 μM PS-liposomes.

**Fig. 3.**
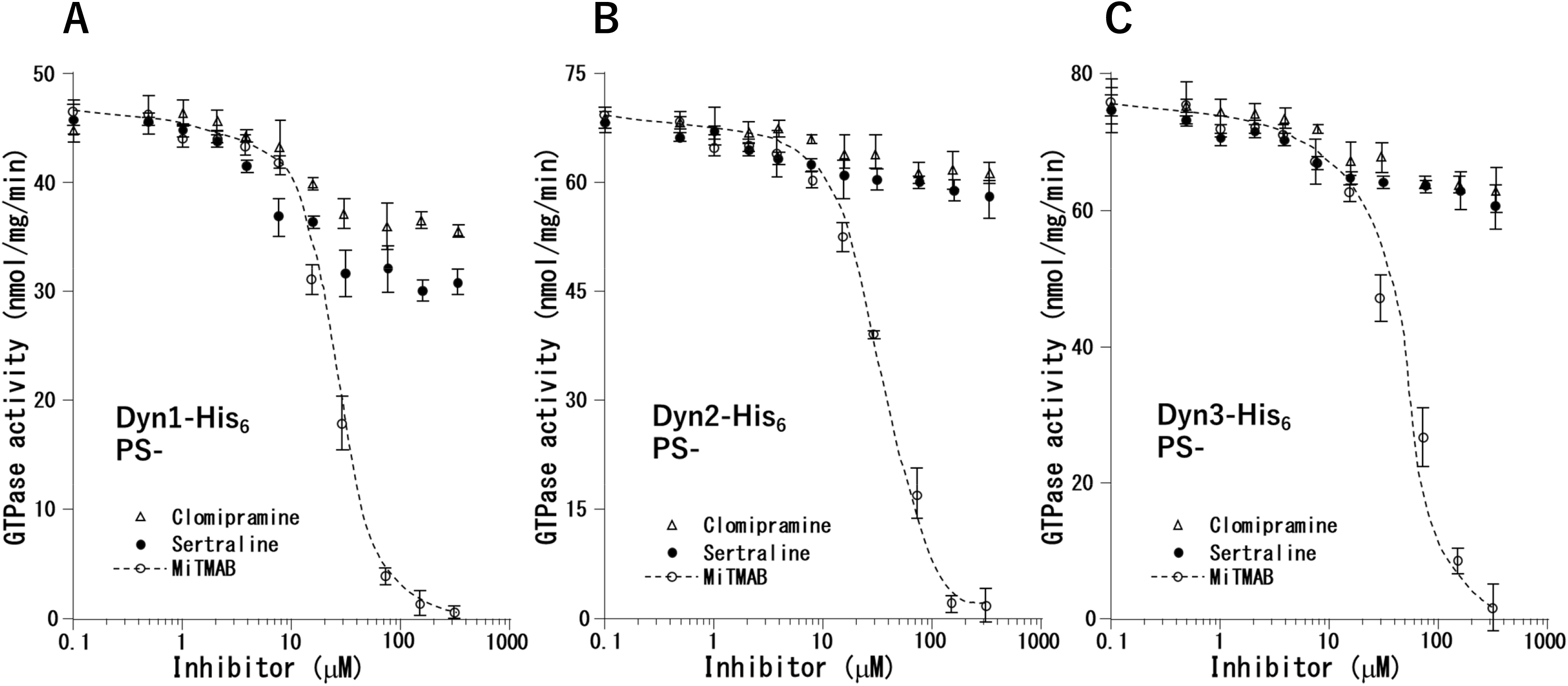
Clomipramine does not inhibit basal intrinsic dynamin GTPase activity. The GTPase activity of purified Dyns-His_6_ (20 nM) was determined in the presence of various concentrations of MiTMAB (hollow circles), sertraline (solid circles), or clomipramine (triangles). All results are representative of at least 3 independent experiments. (A) GTPase activity of Dyn1-His_6_ in the absence of PS-liposomes. (B) GTPase activity of Dyn2-His_6_ in the absence of PS-liposomes. (C) GTPase activity of Dyn3-His_6_ in the absence of PS-liposomes.

### Clomipramine did not inhibit the oligomer formation of Dyn2-His_6_

Dyn GTPase activity is increased by its adoption of helical oligomerization (*18*). BN-PAGE was performed to analyze the effect of clomipramine on the oligomerization of Dyn2-His_6_ (Fig. 4). In the presence of 1% SDS, a denaturant, a single band of approximately 140 kDa, corresponding to the Dyn2-His_6_ monomer, was observed for Dyn2-His_6_ with or without clomipramine treatment, (Fig. 4 lanes 2 and 4, Arrow I). Several bands corresponding to Dyn2-His_6_ oligomers were observed at approximately 300 kDa (Arrow II) and approximately 600 kDa (Arrow III) after BN-PAGE of Dyn2-His_6_ treated with/without clomipramine (Fig. 4 lanes 1 and 3). Moreover, those lanes showed similar band profiles in the higher molecular weight area than at Arrow III (Fig. 4 lanes 1 *versus* 3).

**Fig. 4.**
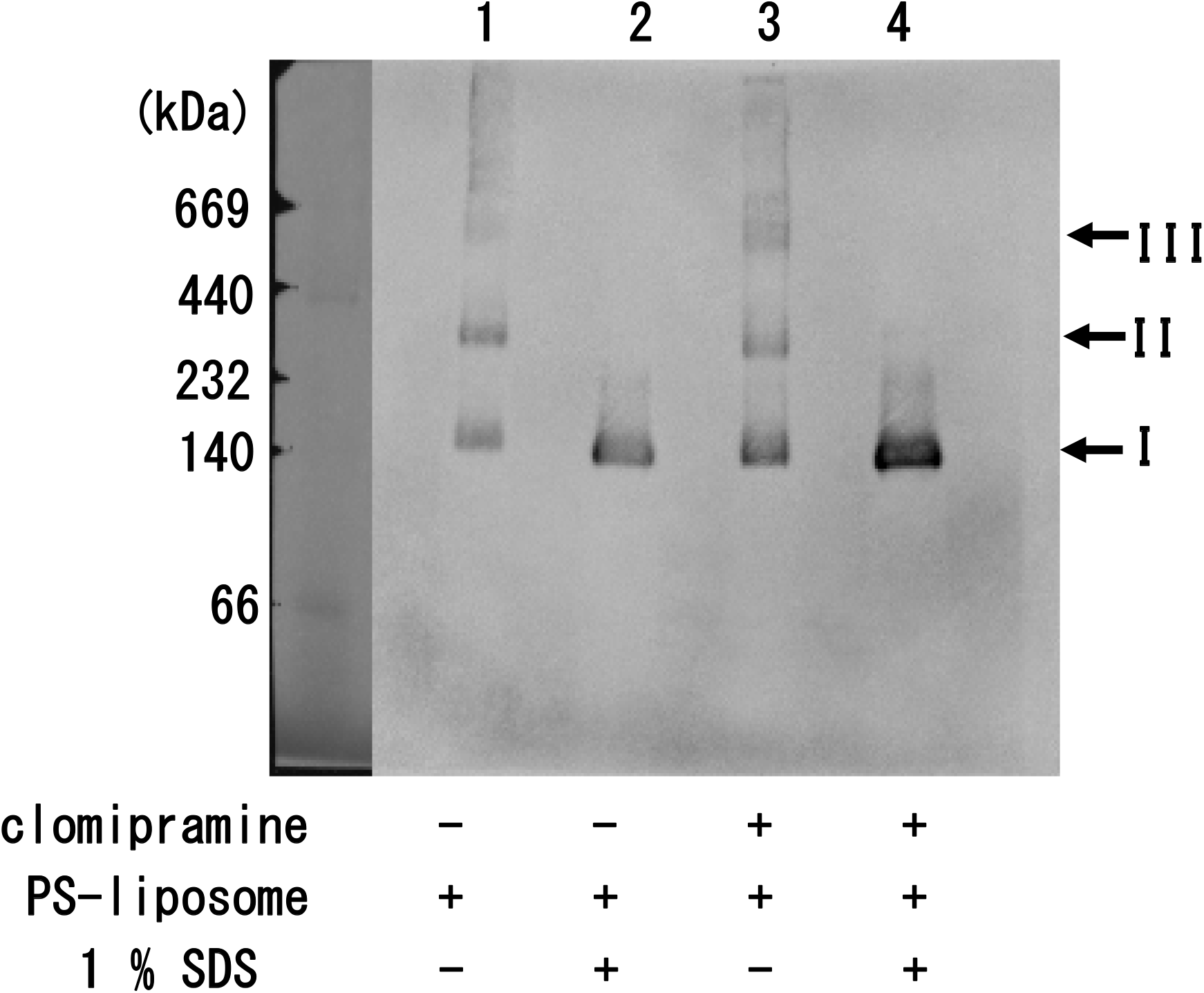
Clomipramine did not affect the oligomerization of Dyn2-His_6_. The effect of clomipramine on the oligomerization of Dyn2-His_6_ was analyzed by BN-PAGE. Lane 1, Dyn2-His_6_ + PS-liposomes; Lane 2, Dyn2-His_6_ + PS-liposomes + 1% SDS; Lane 3, Dyn2-His_6_ + PS-liposomes + clomipramine; Lane 4, Dyn2-His_6_ + PS-liposomes + clomipramine + 1% SDS. Arrow I corresponds to a dynamin monomer. Arrows II and III correspond to dynamin oligomers.

## Discussion

PS-liposomes had been previously reported to stimulate the GTPase activity of Dyn (*19*). As shown in Fig. 1C, PS-liposomes stimulated all purified Dyns-His_6_ GTPase activities in this study. This meant that these purified Dyns-His_6_ can be used to assess the inhibition by clomipramine with GTPase assays. In fact, The IC_50_ values of inhibitors (sertraline, MiTMAB, and clomipramine) for Dyn1-His_6_ obtained from Fig. 2A were not significantly different from those (sertraline; 7.3 ± 1.0 μM, MiTMAB; 24.1 ± 9.4 μM, and clomipramine; 29.9 ± 5.9 μM) obtained in our previous study (*15*).

Clomipramine exhibited inhibition of Dyn2 and Dyn3 GTPase activity similar to its effect on Dyn1 GTPase activity in the presence of PS-liposomes (Fig. 2). Although Dyns-His_6_ GTPase activity in the presence of PS-liposomes was completely lost after 320 μM MiTMAB treatment, Dyns-His_6_ GTPase activity in the addition of clomipramine and sertraline (320 μM) was incompletely lost (Fig. 2A-C). The residual activity is thought to result from GTPase activity in the absence of phospholipids, that is, the so-called basal intrinsic GTPase activity. Thus, these observations show the observation that clomipramine did not strongly inhibit basal intrinsic GTPase activity (Fig. 3A-C). Indeed, the residual activity data between the results without PS addition and with PS addition in the presence of clomipramine were nearly comparable (Fig. 2A *versus* Fig. 3A, Fig. 2B *versus* Fig. 3B, and Fig. 2C *versus* Fig. 3C at 320 μM).

Basal intrinsic GTPase activity tends to decrease after the disassembly of the dynamin oligomer (*20*). Therefore, it is possible that clomipramine does not affect the oligomerization of Dyns-His_6_. Accordingly, we performed BN-PAGE to examine whether clomipramine affects the stabilization of the Dyn2-His_6_ oligomer. The addition of clomipramine did not affect any bands corresponding to Dyns-His_6_ oligomers (Fig. 4, lane 1 *versus* lane 3). Taken together, these results demonstrate that inhibition of Dyns-His_6_ activity did not result from clomipramine-induced destabilization of the oligomer. It is known that the pleckstrin homology domain on dynamin binds to lipids (*21*). Consequently, clomipramine appears to directly interact pleckstrin homology domain, which appears to influence the interaction between PS-liposome and dynamin.

Dyn plays a critical role in clathrin-dependent endocytosis, and SARS-CoV-2 enters into host cells *via* clathrin-dependent endocytosis (*22*). Kato *et al*. mentioned that the tricyclic antidepressant clomipramine suppressed ACE2-mediated SARS-CoV-2 entry (*13, 23*). In this study, we found that clomipramine inhibited the PS-liposome-stimulated GTPase activity of Dyn 2, which was ubiquitously expressed, and Dyn 3, which was expressed in the lung (*4*). Taken together, inhibition of PS-liposome-stimulated GTPase activity by clomipramine may participate in the suppression of ACE2-mediated SARS-CoV-2 entry.

Kato *et al*. also mentioned that the inhibition mechanism by which clomipramine inhibited the entry of SARS-CoV-2 differed from that of other antiviral drugs, such as remdesivir, and suggested the potential for the concomitant use of clomipramine and drugs with other mechanistic effects for treating COVID-19 patients (*23*). Our results reinforce the potential for the concomitant use of these drugs.

## Abbreviations

SARS-CoV-2: severe acute respiratory syndrome coronavirus 2
Dyn: dynamin
MiTMAB: myristyl trimethyl ammonium bromide
PS: L-α-phosphatidyl-L-serine
ACE2: angiotensin-converting enzyme 2
His: histidine
Pi: orthophosphate

## Funding

This work was supported in part by Grants-in-Aid for Scientific Research from the Ministry of Education, Science, Sports and Culture of Japan to KT (21791152) and HM (25430118).

## Author contributions

HM contributed to data analysis and manuscript writing. MO and KT contributed to the collection of data and data analysis. All authors have read and approved the final manuscript.

## Data availability

The data underlying this article are available in the article.

## Conflict of interest

The authors declare that they have no conflict of interest.

## Notes

### Competing Interest Statement

The authors have declared no competing interest.

### Summary of Updates

Section on discussion updated to clarify result; Figure 1, 2, and 3 revised.

## References

1. Moreno-Layseca, P., Jäntti, N.Z., Godbole, R., Sommer, C., Jacquemet, G., Al-Akhrass, H., Conway, J.R.W., Kronqvist, P., Kallionpää, R.E., Oliveira-Ferrer, L., Cervero, P., Linder, S., Aepfelbacher, M., Zauber, H., Rae, J., Parton, R.G., Disanza, A., Scita, G., Mayor, S., Selbach, M., Veltel, S., and Ivaska, J. (2021). Cargo-specific recruitment in clathrin- and dynamin-independent endocytosis. Nature Cell Biology. 23, 1073–1084.

2. Weng, L., Enomoto, A., Miyoshi, H., Takahashi, K., Asai, N., Morone, N., Jiang, P., An, J., Kato, T., Kuroda, K., Watanabe, T., Asai, M., Ishida-Takagishi, M., Murakumo, Y., Nakashima, H., Kaibuchi, K., and Takahashi, M. (2014). Regulation of cargo-selective endocytosis by dynamin 2 GTPase-activating protein girdin. The EMBO journal. 33, 2098–2112.

3 Henley, J.R., Krueger, E.W., Oswald, B.J. and McNiven, M.A. (1998). Dynamin-mediated internalization of caveolae. The Journal of cell biology. 141, 85–99.

4. Reis, C.R., Chen, P.H., Srinivasan, S., Aguet, F., Mettlen, M. and Schmid, S.L. (2015). Crosstalk between Akt/GSK3β signaling and dynamin-1 regulates clathrin-mediated endocytosis. The EMBO journal. 34, 2132–46.

5. Ramachandran, R. and Schmid, S.L. (2018). The dynamin superfamily. Current Biology 28, R411–R416.

6. van Dam, E.M. and Stoorvogel, W. (2002). Dynamin-dependent transferrin receptor recycling by endosome-derived clathrin-coated vesicles. Molecular biology of the cell. 13, 169–82.

7. Muhlberg, A.B., Warnock, D.E. and Schmid, S.L. (1997). Domain structure and intramolecular regulation of dynamin GTPase. The EMBO journal. 16, 6676–83.

8. Bashkirov, P.V., Akimov, S.A., Evseev, A.I., Schmid, S.L., Zimmerberg, J. and Frolov, V.A. (2008). GTPase Cycle of Dynamin Is Coupled to Membrane Squeeze and Release, Leading to Spontaneous Fission. Cell. 135, 1276–1286.

9. Quan, A., McGeachie, A.B., Keating, D.J., van Dam, E.M., Rusak, J., Chau, N., Malladi, C.S., Chen, C., McCluskey, A., Cousin, M.A., and Robinson, P.J. (2007). Myristyl Trimethyl Ammonium Bromide and Octadecyl Trimethyl Ammonium Bromide Are Surface-Active Small Molecule Dynamin Inhibitors that Block Endocytosis Mediated by Dynamin I or Dynamin II. Molecular Pharmacology. 72, 1425–1439.

10. Helle, F. and Dubuisson, J. (2007). Hepatitis C virus entry into host cells. Cellular and Molecular Life Sciences. 65, 100.

11. Li, W., Moore, M.J., Vasilieva, N., Sui, J., Wong, S.K., Berne, M.A., Somasundaran, M., Sullivan, J.L., Luzuriaga, K., Greenough, T.C., Choe, H., and Farzan, M. (2003). Angiotensin-converting enzyme 2 is a functional receptor for the SARS coronavirus. Nature. 426, 450–4.

12. Hoffmann, M., Kleine-Weber, H., Schroeder, S., Krüger, N., Herrler, T., Erichsen, S., Schiergens, T.S., Herrler, G., Wu, N.H., Nitsche, A., Müller, M.A., Drosten, C., and Pöhlmann, S. (2020). SARS-CoV-2 Cell Entry Depends on ACE2 and TMPRSS2 and Is Blocked by a Clinically Proven Protease Inhibitor. Cell. 181, 271–280 e8.

13. Kato, Y., Yamada, S., Nishiyama, K., Satsuka, A., Re, S., Tomokiyo, D., Lee, J.M., Tanaka, T., Nishimura, A., Yonemitsu, K., Asakura, H., Ibuki, Y., Imai, Y., Kamiya, N., Mizuguchi, K., Kusakabe, T., Kanda, Y., and Nishida, M. (2021). Clomipramine suppresses ACE2-mediated SARS-CoV-2 entry. bioRxiv, 2021.03.13.435221.

14. Vercauteren, D., Vandenbroucke, R.E., Jones, A.T., Rejman, J., Demeester, J., De Smedt, S.C., Sanders, N.N. and Braeckmans, K. (2010). The use of inhibitors to study endocytic pathways of gene carriers: optimization and pitfalls. Molecular therapy: the journal of the American Society of Gene Therapy. 18, 561–9.

15. Otomo, M., Takahashi, K., Miyoshi, H., Osada, K., Nakashima, H. and Yamaguchi, N. (2008). Some Selective Serotonin Reuptake Inhibitors Inhibit Dynamin I Guanosine Triphosphatase (GTPase). Biological and Pharmaceutical Bulletin. 31, 1489–1495.

16. Takahashi, K., Miyoshi, H., Otomo, M., Osada, K., Yamaguchi, N. and Nakashima, H. (2010). Suppression of dynamin GTPase activity by sertraline leads to inhibition of dynamin-dependent endocytosis. Biochemical and biophysical research communications. 391, 382–7.

17 Takahashi, K., Otomo, M., Yamaguchi, N., Nakashima, H. and Miyoshi, H. (2012). Replacement of Arg-386 with Gly in Dynamin 1 Middle Domain Reduced GTPase Activity and Oligomer Stability in the Absence of Lipids. Bioscience, Biotechnology, and Biochemistry. 76, 2195–2200.

18 Stowell, M.H.B., Marks, B., Wigge, P. and McMahon, H.T. (1999). Nucleotide-dependent conformational changes in dynamin: evidence for a mechanochemical molecular spring. Nature Cell Biology. 1, 27–32.

19. Tuma, P.L., Stachniak, M.C. and Collins, C.A. (1993). Activation of dynamin GTPase by acidic phospholipids and endogenous rat brain vesicles. Journal of Biological Chemistry. 268, 17240–17246.

20 Warnock, D.E., Baba, T. and Schmid, S.L. (1997). Ubiquitously expressed dynamin-II has a higher intrinsic GTPase activity and a greater propensity for self-assembly than neuronal dynamin-I. Molecular biology of the cell. 8, 2553–2562.

21 Vallis, Y., Wigge, P., Marks, B., Evans, P.R. and McMahon, H.T. (1999). Importance of the pleckstrin homology domain of dynamin in clathrin-mediated endocytosis. Current Biology. 9, 257–263.

22 Yang, H., Yuan, H., Zhao, X., Xun, M., Guo, S., Wang, N., Liu, B. and Wang, H. (2022). Cytoplasmic domain and enzymatic activity of ACE2 are not required for PI4KB dependent endocytosis entry of SARS-CoV-2 into host cells. Virologica Sinica. 37, 380–389.

23 Kato, Y., Nishiyama, K., Nishimura, A., Noda, T., Okabe, K., Kusakabe, T., Kanda, Y. and Nishida, M. (2022). Drug repurposing for the treatment of COVID-19. Journal of Pharmacological Sciences. 149, 108–114.

